# Sparse RNNs can support high-capacity classification

**DOI:** 10.1101/2022.05.18.492540

**Authors:** Denis Turcu, L. F. Abbott

## Abstract

Feedforward network models performing classification tasks rely on highly convergent output units that collect the information passed on by preceding layers. Although convergent output-unit like neurons may exist in some biological neural circuits, notably the cerebellar cortex, neocortical circuits do not exhibit any obvious candidates for this role; instead they are highly recurrent. We investigate whether a sparsely connected recurrent neural network (RNN) can perform classification in a distributed manner without ever bringing all of the relevant information to a single convergence site. Our model is based on a sparse RNN that performs classification dynamically. Specifically, the interconnections of the RNN are trained to resonantly amplify the magnitude of responses to some external inputs but not others. The amplified and non-amplified responses then form the basis for binary classification. Furthermore, the network acts as an evidence accumulator and maintains its decision even after the input is turned off. Despite highly sparse connectivity, learned recurrent connections allow input information to flow to every neuron of the RNN, providing the basis for distributed computation. In this arrangement, the minimum number of synapses per neuron required to reach maximum memory capacity scales only logarithmically with network size. The model is robust to various types of noise, works with different activation and loss functions and with both backpropagation- and Hebbian-based learning rules. The RNN can also be constructed with a split excitation-inhibition architecture with little reduction in performance.

## Introduction

Binary classification is a basic task that involves dividing stimuli into two groups. Machine-learning solutions to this task typically use single- or multi-layer perceptrons [1] in which, almost invariably (but see [2]), the output that delivers the network’s decision comes from a unit that collects information from all of the units in the previous layer. Collecting all of the evidence in one place (i.e. in one unit) is an essential element in the design of these networks. In humans and other mammals, tasks like this are likely performed by neocortical circuits that have a recurrent rather than feedforward architecture, and where there are no obvious highly convergent ‘output’ neurons. Instead, all of the principal neurons are sparsely connected to a roughly equal degree. This raises the question of whether a network can classify effectively if the information needed for the classification remains dispersed across the network rather than being concentrated at a single site. Here we explore how and how well recurrent networks with sparse connections and no convergent output unit can perform binary classification.

We study sparse RNNs that reach decisions dynamically. Despite their sparse connectivity, these networks are able to compute distributively by propagating information across their units. To add biological realism, we also constrain our sparse RNN to have a split excitation-inhibition architecture. The model maintains high performance despite this constraint. To investigate capacity and accuracy, networks were trained by back-propagation through time (BPTT). With extensive training, these models can categorize up to two input patterns per plastic synapse, matching the proven limit of the perceptron [3]. The model is robust to different types of noise, training methods and activation functions. The number of recurrent connections per neuron needed to reach high performance scales only logarithmically with network size. To investigate biologically plausible learning, we also constructed networks using both one-shot and iterative Hebbian plasticity. Although performance is significantly reduced compared to BPTT, capacity is still proportional to the number of plastic synapses in the RNN.

## Results

### The Model

We built a sparse RNN [Figure 1A] and evaluated its performance on a typical binary classification task. The RNN consists of *N* units described by a vector **x** that evolves in time according to

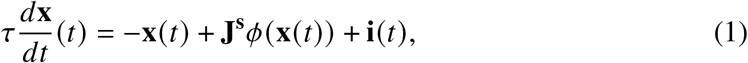

where *τ* is a time constant and **i** is the external input being classified. The response function *ϕ*(·) is, in general, nonlinear, monotonic, non-negative and bounded; we use either a shifted, scaled and rectified hyperbolic tangent (Methods) or a squared rectified linear function [Figure 1B]. Both response functions performed well; we use the modified hyperbolic tangent for all the results we report. Importantly, the connection matrix **J**^**s**^ is sparse with only *f N*^2^ randomly placed non-zero elements, for 0 < *f* < 1. Sparsity is enforced by a mask that also constrains the sparseness during training Methods.

**Figure 1:**
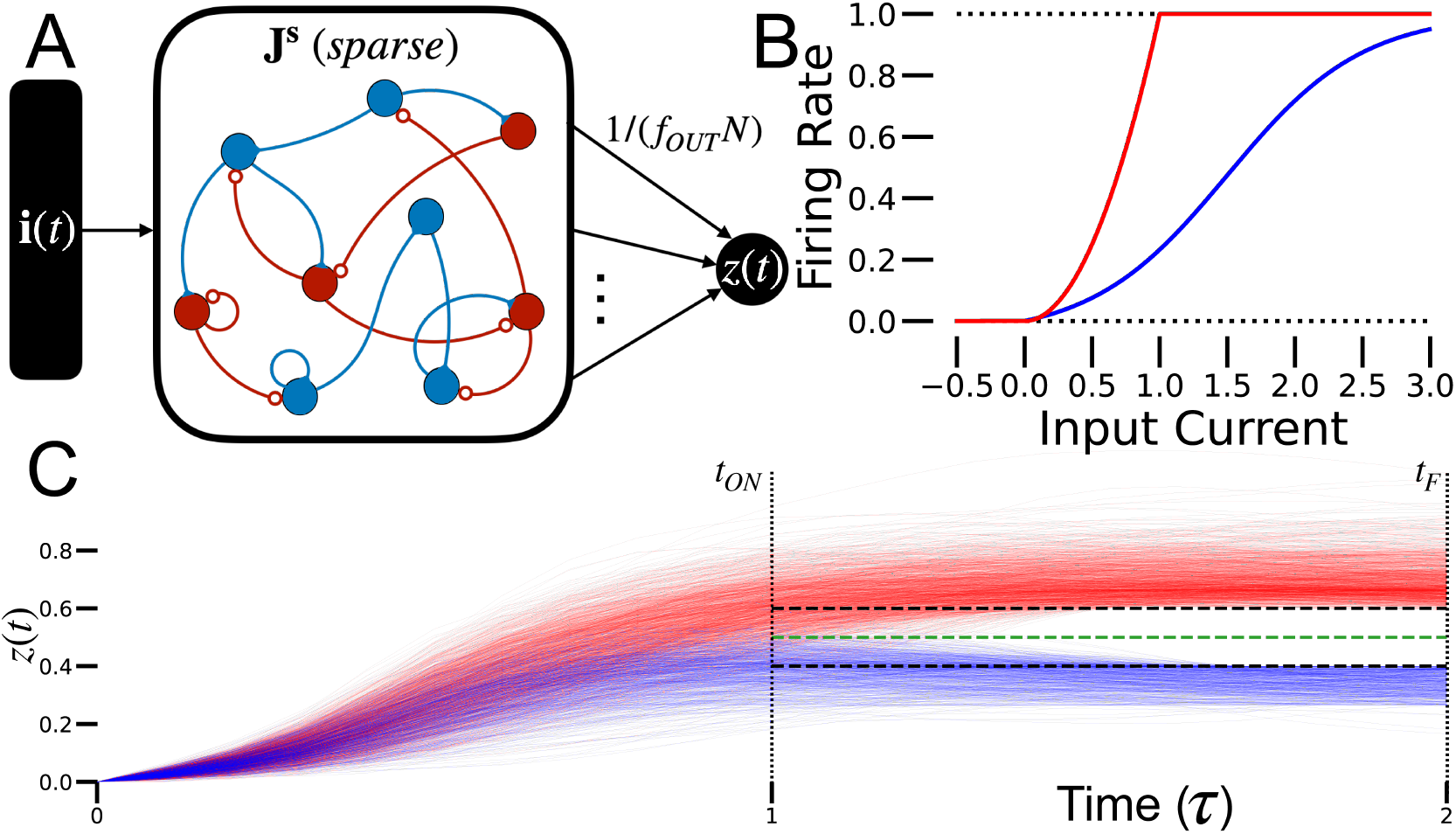
**(A)** The sparse RNN. Recurrent connections are plastic, but input and output connections are fixed and uniform. We consider both mixed and split excitation-inhibition networks (the split architecture is shown here). **(B)** Examples of activation functions used. Red line is squared rectified linear function, bounded by 1. Blue line is a shifted, scaled and rectified hyperbolic tangent function. **(C)** Illustration of target dynamics training method. Readout activity of trained RNN is shown in red for +1-labeled inputs and in blue for −1-labeled inputs. The corresponding targets *T_+_*and *T*_ are shown as dashed black lines. The threshold *θ* is the dashed green line.

Categorization requires the RNN to correctly match inputs with associated binary labels. Inputs are *N* – dimensional vectors, called patterns, with each component chosen randomly and independently from a uniform distribution between -1 and 1. The patterns are represented by *P N* -component vectors *ξ* ^*μ*^ with elements 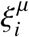 for *i* = 1, 2, …, *N* and *μ* = 1, 2, …, *P*. The label for pattern *μ, q*^*μ*^, is chosen randomly for each pattern to be either -1 or 1, with equal probability. The category assigned by the RNN is determined by averaging the activity of *f*_out_*N* randomly chosen units in the RNN, a quantity denoted by *𝓏(t)* [Figure 1A]. We typically set *f*_out_ = *f*, except in Figure 1B. The sparse average over neurons, *𝓏(t)*, measured at a specified time *t*_OUT_, reports the decision of the network. Specifically, the category determined by the RNN is defined as -1 if *𝓏(t*_OUT_) is less than a threshold *θ*, and 1 otherwise. The fraction of input-label pairs that are categorized correctly quantifies the accuracy of the sparse RNN, and its capacity is the number of patterns that can be categorized to a given level of accuracy.

Each categorization run starts at time 0 with the initial RNN activity set to zero, **x**(0) = **0**, and **i** set to one of the input patterns, **i** = *ξ* ^*μ*^ for a randomly chosen *μ* value. The input remains on at a constant level for a duration *t*_ON_ and then **i** is set to zero. The trial terminates at time *t*_F_. After training, the network’s decision can be read out at any time *t*_ON_ < *t*_OUT_ <*t*_F_ with minimal effect on performance.

Training was used to adjust only the recurrent connections; the input connections are one-to-one with unit weights and the output connections are sparse and fixed at 0 or 1 /(*f*_OUT_*N)* [Figure 1A]. Training did not include noise in the sparse RNN dynamics, but two types of noise were incorporated for testing (see below). For our initial studies, the weights of the connections, given by **J**^**s**^, were set by backpropagation through time (BPTT), with the goal of generating the correct associated label for each input pattern. This training method can incorporate the sparseness constraints imposed on **J**^**s**^ and, as discussed later, can also impose a split excitation-inhibition architecture (Methods).

To train the network, label-specific target dynamics were imposed on the output *𝓏(t)*, at certain times [Figure 1C]. The target dynamics consist of constant target values *T*_+(−)_ (dashed black curves in [Figure 1C]). The target values for the ± 1 categories satisfy *T*_+_>*T*_−_ for all times. The loss function penalizes the RNN at all times *t*_ON_ <*t* < *t*_F_ when the output is below (+1 category) or above (−1 category) the target in proportion to the absolute difference between *𝓏(t)*, and *T*_+(−)_. This loss function implicitly defines a discrimination threshold 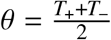 (dashed green curve in [Figure 1C]) for computing the decision (Methods). The targets were chosen so that the threshold was small, 0.1 ≤ *θ* ≤ 0.6, to take advantage of the supralinear behavior of the neural response function near 0, which enhanced performance.

The targets and threshold were constant in time to allow the network to act as an evidence accumulator.

Training length varied depending on the convergence criteria and to maintain reasonable running times. We ran 5000 BPTT epochs for results in [Figure 2D], and typically at most 500 epochs for all other results. We found that performance improvement was minimal beyond 500 epochs, especially for small networks, and in some cases, particularly in which fewer pattern-label pairs were shown, 300, 200 or even 150 epochs sufficed to reach good performance, demonstrate the results and gain negligible improvement from epoch to epoch.

**Figure 2:**
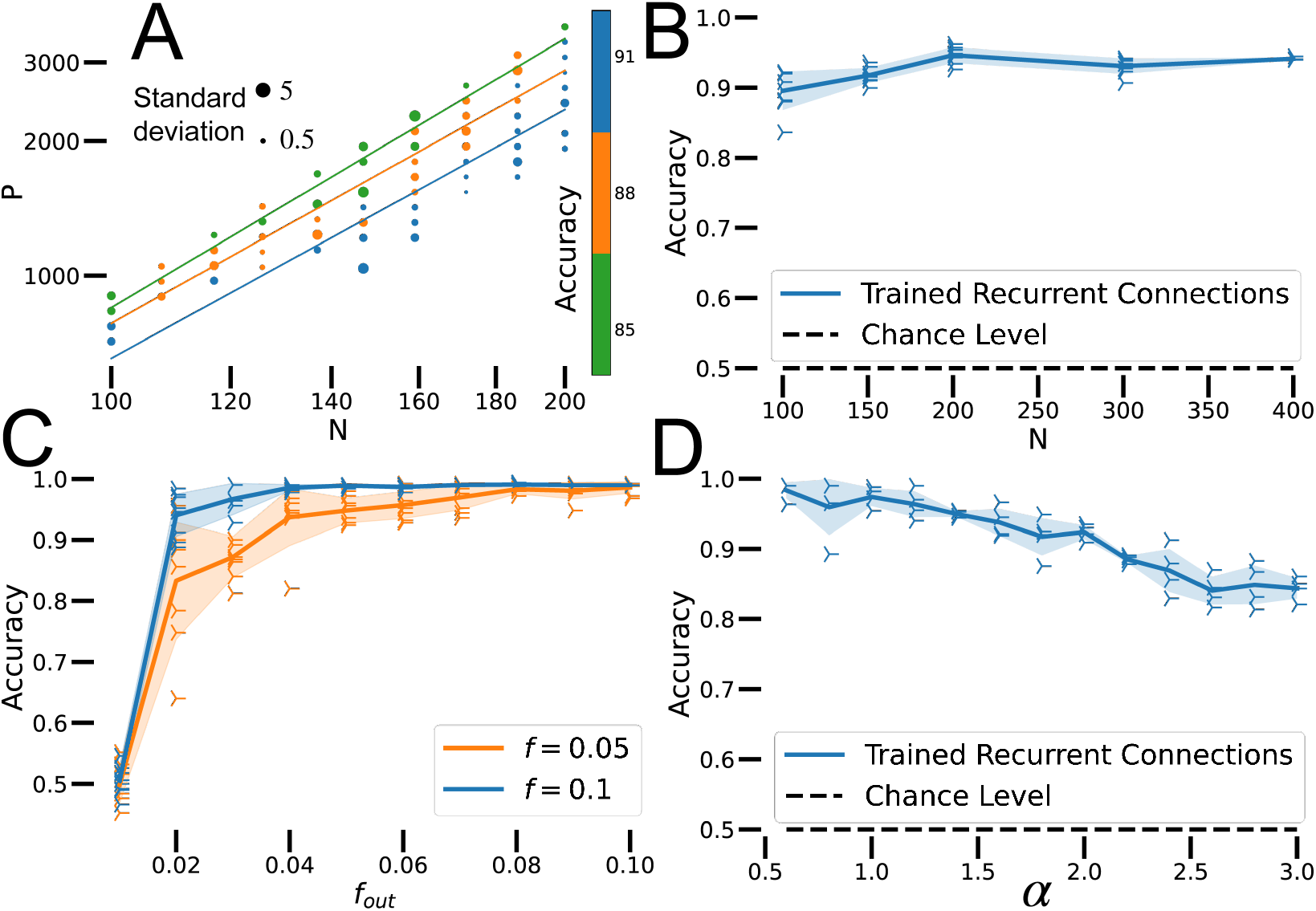
**(A)** Accuracy for various combinations of *N* and *P*. Colors represent accuracy, grouped in bins spanning 3%. Lines are fitted to every group of binned accuracy values. *f* = 0.1, # epochs = 200, *n* = 4. **(B)** Binary classification performance of sparse RNNs of various sizes. Each RNN classifies *P* = *α f N* ^2^ inputs. = 0.1, *α* = 1, # epochs = 500, *n* = 4. **(C)** Sparse RNN performance as a function of the readout sparsity for RNNs with two different levels of recurrent sparsity. *N* = 100, *α* = 0.5, # epochs = 500,*n* = 6. **(D)** Accuracy of sparse RNNs as a function of *α* = *P*/(*f N*^2^) Runs were truncated at 99% accuracy. *N* = 100, *f* = 0.05, # epochs = 5000, *n* = 4.

RNNs can be trained to perform categorization across a range of the temporal parameters *t*_ON_, *t*_OUT_ and *t*_F_. Networks performed well for trial durations in the range *τ* < *t*_F_ < 15τ with *t*_ON_ = *t*_F_ / 2 when sampled within the range 3 *t*_F_ / 4 to *t*_F_. Performance was poor when the trial duration was very small, i.e. *t*_F_ < 0.2*τ*. For the results reported below, we set *t*_ON_ = *τ, t*_F_ = 2*τ* and sampled the RNN with 1.5*τ* < *t*_OUT_ < 2*τ*.

Note that the readout is not learned and is as sparsely connected as the units of the RNN. Thus there is no special convergence of information onto the read out. In addition, each RNN unit receives only one component of the input vector and units are sparsely interconnected.

Thus, there is no locus where information converges. Instead, the network units must solve the classification task collectively.

### Network Performance

Sparse RNNs trained with BPTT can memorize a number of pattern-label pairs proportional to the number of plastic synapses, *P* = *α f N* ^2^ [Figure 2A]. To verify this, we determined the slopes of lines of constant performance on a plot of log *P* versus log *N* [Figure 2A]. To avoid excessive training time, all networks in Figure 2A were trained for 200 epochs. Even with limited training, the *R*^2^ of the regressions is 0.95 ± 0.03 and the slope is 1.91 ± 0.07. We observed that training converges faster for smaller networks, which could affect the comparison of networks across many sizes. Removing networks smaller than 115 neurons from this analysis yielded a similar regressions value, *R*^2^ = 0.93 ± 0.03 and a slope of 2.08 ± 0.05. These results support that *P ∼N*^2^, that is, sufficiently large sparse RNNs can memorize a number of pattern-label pairs proportional to the square of their neuron numbers or to the first power of their plastic synapse counts.

For given values of and *f*, the categorization accuracy for *P* = *α f N* ^2^ patterns is independent of network size, for large *N* [Figure 2B]. Recall that the network output is obtained by averaging the activity of *f*_out_*N* units. This level of output sparsity does not impair performance as long as *f*_out_ is comparable to *f* [Figure 2C]. We use *f*_out_ = *f* for all further results.

Well trained sparse RNNs reach high memory capacity and do not manifest “blackout catastrophe”. Often they can perfectly memorize one input-label pair for each plastic synapse (*α* = 1) if trained for a sufficient time. Alternatively, they can classify up to two patterns for each plastic synapse with accuracy > 90% [Figure 2D]. Accuracy decreases as α increases, but the decrease in accuracy is gradual, even for *α* > 2. Thus, sparse RNNs do not completely forget previously learned patterns when pushed above a critical value of *α*. Instead, their performance decreases gradually for increasing numbers of patterns.

### Minimum spareseness

Sparse RNNs categorize by distributing the computation across all the elements of the network. Because all the information is not brought together at a single locus, that information must flow freely through the network to generate a correct categorization. The importance of information propagation is evidenced by the reduced performance of RNNs at extreme sparsity levels [Figure 3A].

**Figure 3:**
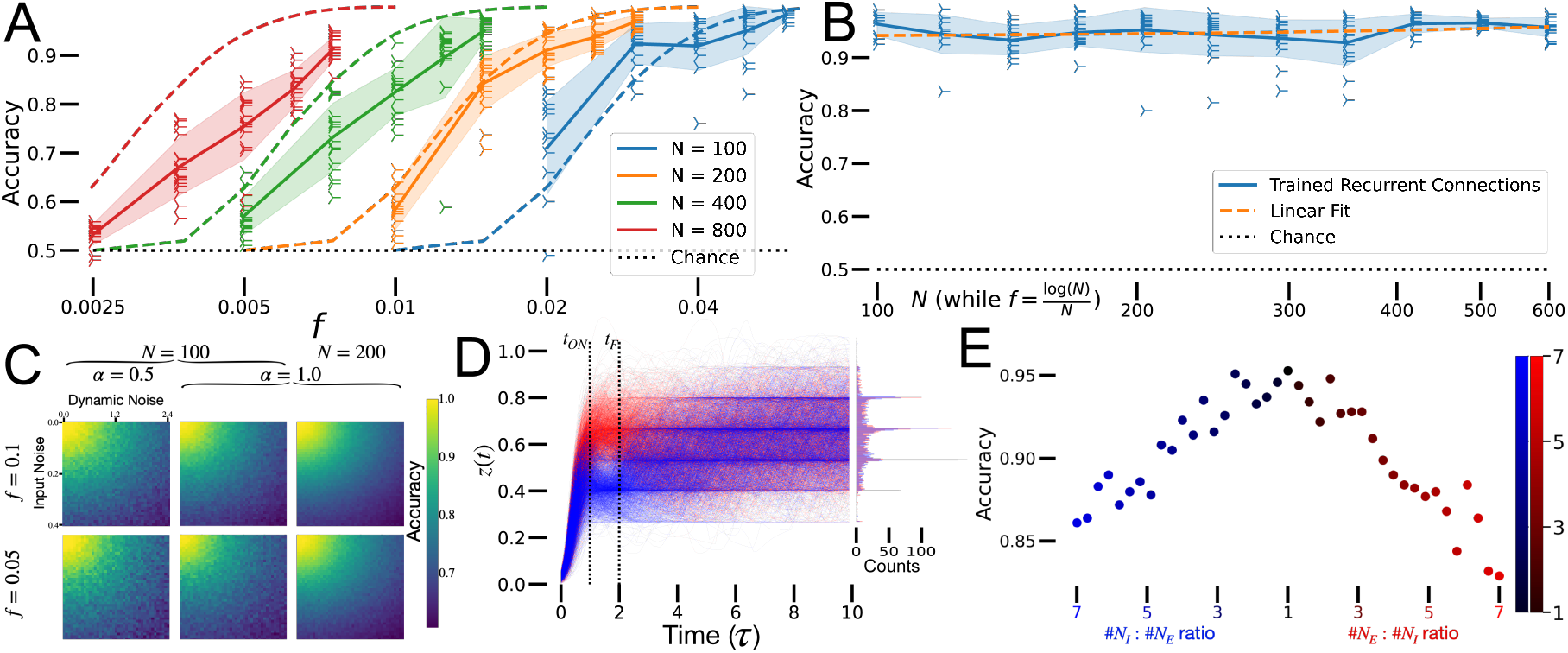
**(A)** Performance of sparse RNNs at extreme sparsity levels as a function of model size (scatter points; solid lines are averages and shaded areas are standard deviations). Predictions based on empirical analysis of random directed graphs (dashed lines). *α* = 0.5, # epochs = 500,*n* = 16. **(B)** Performance of nearly fully connected sparse RNNs at the extreme sparsity level *f* = log *(N)/N* for various networks sizes – the size of the strongly connected component is at least 0.95 *N* (blue scatter points; blue solid lines are averages and blue shaded areas are standard deviations). Linear fit of all the points is shown as orange dashed line. *α* = 0.5, # epochs = 1000,*n* = 16. **(C)** Noise robustness heat maps for sparse RNNs. The horizontal axis indicates the level of dynamic noise, defined as the standard deviation of the dynamic noise divided by the mean of the activity in the RNN, and the vertical axis indicates the standard deviation of input noise. Sparse RNNs maintain high accuracy over a wide range of noises levels. # epochs = 150. **(D)** Output of an example sparse RNN over time. Model is trained for 2*τ*, and input is on for *τ* .*N* = 300, *f* = 0.05, *α* = 1, # epochs = 500. **(E)** Performance of the E/I sparse RNNs as a function of the E/I ratio. *N* = *N*_*E*_ +*N*_*I*_ = 100, *f* = 0.1, *α* = 1, # epochs = 200.

Signal propagation in networks can be characterized by their adjacency graphs. Randomly generated undirected graphs are disconnected at extreme sparsity levels according to the Erdös-Rényi model [4]. However, as in biological networks, our sparse RNNs are based on directed graphs, meaning we cannot fully apply the analytic results developed by [4]. A directed graph of size *N* with connection sparsity *f* consists of a large strongly connected sub-graph of size *N* − *k* and *k* other nodes with some probability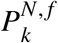 (Methods). We assumed the main sub-graph of size *N − k* performs the task on all *αfN*^2^ inputs without help or interference from the other *k* nodes, and computed the expected accuracy of the sparse RNNs estimating 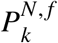 empirically. We also assumed that the network correctly classifies *αf*(*N* −)^2^(*N* − *k*) / *N* inputs and performs at chance level on the rest, and we computed the estimated accuracy of a network of size *N* with only *N* − *k* functional units, 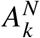. Thus, the expected accuracy of the sparse RNN with *N* neurons and sparsity 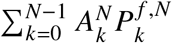, shown as dashed lines in Figure 3A.

The performance of our models roughly fits the predictions based on the empirical analysis of random directed graphs [Figure 3A]. In particular, the shape of the performance drop qualitatively matches our expectations and the location of the initiation of the performance drop appears to shift logarithmically with network size, as expected from our empirical analysis of directed graphs. Discrepancies between simulations and empirical analysis predictions are apparent for large network sizes. One reason for the increasing discrepancy with network size may lie in the difficulty of training extremely sparse large networks, which require more training epochs. Running time per epoch scales like *N*^4^ in our task, since both the number of plastic synapses and the number of inputs scale like *N*^2^, making it very difficult to train large networks for many more epochs. Another reason for the discrepancy may lie in our assumptions in computing 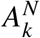. We assumed that the *k* separated nodes would not interfere with network decisions. Though we accounted for the likelihood that not all *f N* readout neurons will be part of the main connected sub-graph through the (*N* − *k*)/*N* factor from 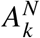, we suspect that reading out from one of the *k* separated neurons introduces further noise and impairs performance. This effect is intensified at extreme sparsity values, where large networks still perform above chance level.

This evidence suggests that it is sufficient for our sparse RNNs to have a strongly connected adjacency graph to perform well. For undirected graphs, Erdös-Rényi results provide a basis for analytically computing the number of connections each node needs for a graph to be connected with probability near 1 [4]. Specifically, a random graph of size *N* with *f N*^2^ connections is fully connected with probability exp (− exp (− 2 *f N +* log *N))*, in the limit of large graph sizes. If we set that probability to 1 − *ε* for *ε* << 1, the average number of synapses per neuron is *f N*^2^/ *N* = *f N* = log *(N)/*2 − log [log (1 / (1 − *ε*)], which scales as log *N* for fixed *ε*. Moreover, the distribution of the number of synapses per neuron follows a binomial distribution, according to the definition of the adjacency graph. For directed graphs, such as our sparse RNN model, we observe the same logarithmic scaling in our empirical analysis of their adjacency graphs.

To test this hypothesis, we simulated sparse RNNs of various sizes *N* and having the extreme sparsity level *f* = log (*N*)/*N*. We generated the sparse RNNs randomly and rejected all networks whose strongly connected component was smaller than 0.95. We found that these networks performed the task well and at a constant level across network size under the same training conditions [Figure 3A]. The line fit to these data points has a slope of 0.00003 ≈ 0. The results summarized in Figures 3A,B] support the need for collective decision making across multiple units.

### The Effects of Noise

We tested the robustness of sparse RNNs to two types of noise, injected after training. ‘Input noise’ is added to the input of the RNN whenever **i**(*t*) ≠ 0. This noise changes for every trial but remains constant once the input is on in each trial. ‘Dynamic noise’ is added to each RNN unit at all times during each trial and changes at every time step. Both noise sources are generated randomly and independently for each unit (and time for dynamic noise) from a Gaussian distribution. Trained sparse RNNs of various sizes, sparsities and capacities maintain high performance over a wide range of noise values [Figure 3C]. Larger sparse RNNs respond to noise more predictably than smaller models (note the smoother heat maps). Sparse RNNs trained to memorize fewer input patterns are more noise resistant (note the yellow color covers a larger area in Figure 3C.

### Output Dynamics

We examined the dynamics of network outputs before, during and after the time of readout [Figure 3D]. BPTT-trained sparse RNNs develop their decisions incrementally in time, acting as evidence accumulators, and they maintain their decisions for a period of time following the readout time. Initially, the models integrate input information while pattern input is present, allowing the sparse RNN to separate into two label-dependent output distributions. After the input is turned off, the model maintains the decision, typically for the same duration as it was trained. During this time, the output distributions remain separated. At longer times, the distributions slowly mix and discrimination between the two categories is lost. The sparse RNNs do not construct or reach two fixed points that correspond to the two labels [5]. Instead, although the activity is sustained for a while, eventually the separability of the input classes is lost [Figure 3D, histogram].

### E-I Architecture

In cortical networks, excitatory and inhibitory neurons separately provide positive and negative inputs to each neuron. A split excitation-inhibition architecture can be enforced on sparse RNNs during training by constraining the connections matrix **J**^**s**^ such that elements in individual columns of the sparse matrix all have the same sign. The E or I identity is assigned randomly for each column, with possible bias towards one of the two identities. In our investigations, the ratio between the number of E neurons,, *N*_*E*_and the number of I neurons, *N*_*I*_, where *N*_*E*_ + *N*_*I*_ = *N*, took a range of values.

Split excitation-inhibition architecture does not significantly impair the performance of sparse RNNs performing classification [Figure 3E]. A balanced ratio of 1 between the number of excitatory and inhibitory neurons works best and achieves performance similar to the unconstrained architecture [compare to Figure 2D]]. Interestingly, a ratio skewed towards more excitatory neurons appears to be more detrimental than a bias in the inhibitory direction.

### Effect of response nonlinearity and loss function

Sparse RNNs can categorize using different activation and loss functions, although certain of these are better than others [Table 1]. The activation functions that provided best results had a slight supralinear behavior for small and positive inputs and were bounded by above and below. The supralinear behavior helps separate the output distributions more rapidly by keeping low activity low while pushing high activity disproportionately higher.

**Table 1:** Accuracy for combinations of response nonlinearities and loss functions. The hyperbolic tangent response non-linearity was shifted, rectified and scaled to 1. The polynomial response nonlinearities were rectified and bounded by 1. The subscript on the threshold loss function indicates the amount of time the network was trained on. Best results are in green and worst in red. *N* = 100, *f* = 0.1, *α* = 1, # epochs = 300, *n*= 8.

| | $\mathcal{L}^{\text{targets}}$ | $\mathcal{L}_{2\tau}^{\text{threshold}}$ | $\mathcal{L}_{\tau}^{\text{threshold}}$ | $\mathcal{L}_{0.1\tau}^{\text{threshold}}$ |
| --- | --- | --- | --- | --- |
| Modified tanh($x$) | **100 ± 0.0** | 97.3 ± 0.4 | 98.2 ± 0.4 | 96.3 ± 0.1 |
| Bounded ReLU($x$) | 99.9 ± 0.1 | 97.9 ± 0.1 | 98.7 ± 0.1 | 94.3 ± 0.7 |
| Bounded ReLU($x^2$) | **100 ± 0.0** | 96.7 ± 0.9 | 97.4 ± 0.8 | 96.3 ± 0.7 |
| Bounded ReLU($x^3$) | 96.9 ± 0.3 | 97.4 ± 0.4 | 95.9 ± 0.5 | **87.3 ± 2.3** |

In addition to the loss function ℒ^targets^ described above, we trained networks with a loss ℒ^threshold^ that depends on the quantity −*q*^*μ*^(*𝓏(t)* − θ) summed from time *t*_F_ − Δ*t*. to time *t*_F_. A nonlinear mapping assured that positive values, i.e. categorization miss-matches, contributed large penalties to the loss, whereas negative values contributed little (Methods). This loss function worked well, but produced accuracy values about 5% worse than the results we report using the previously described loss [Table 1].

### Hebbian Learning

Although it is useful for constructing networks for study, BPTT is not a biologically realistic way to train networks to perform categorization. It is possible to construct sparse RNNs that perform binary classification, albeit with less capacity, using a closed expression for the connection matrix **J**^**s**^. This connection matrix can be interpreted as the result of sequentially applying a Hebbian rule to the connections of the sparse RNN learning the patterns, with one exposure each. For this reason, we called it one-shot (OS). Specifically, we set the elements of **J**^**s**^ to

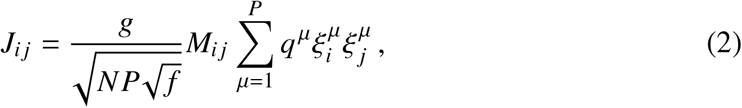

where **M**, with elements equal to either 0 or 1, is the sparsity mask matrix and is set to the value 5.62 to optimize performance. We found this value of *g* by using a grid search and chose the value that performed best.

Sparse RNNs initialized using the OS method memorize a number of pattern-label pairs proportional to the number of plastic synapses, *p* = *α* _OS_ *f N*^2^, similarly to BPTT-trained sparse RNNs, except *α* _OS_ *≈ α* _BPTT_ / 1000 ≈ 0.001 [Figure 4OS]. We show the performance of many sparse RNNs of various sizes being presented different numbers of pattern-label pairs. We divided the achieved accuracy into bins spanning 2% and fit lines to all networks that fall in one bin. The *R*^2^ of the 9 line fits is 0.9936 ± 0.0050. The average slope of those lines is 2.047 ± 0.030. These results suggest that the number of pattern-label pairs memorized up to a given accuracy by the sparse RNNs scales proportionally with the number of plastic synapses, in particular *P* ∼ *N*^2^.

**Figure 4:**
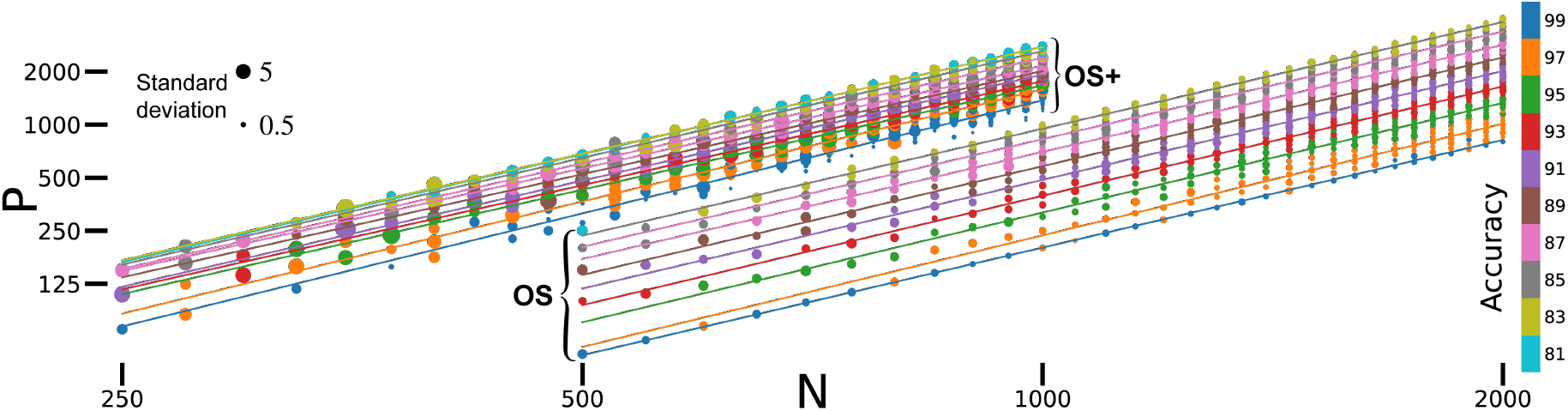
Accuracy for OS and OS+ methods at various combinations of *N* and *P*. Colors represent accuracy, grouped in bins spanning 2%. Lines are fitted to every group of binned accuracies, with all slopes > 2 and regression *r* > 0.98. Size of the scatter points is proportional to the standard deviation (inset). f = 0.1. OS: *n*= 10. OS+:*n* = 5, *ϵ* = 0.02, # epochs = 150.

We also considered an extension from OS to OS+ that includes plasticity of the recurrent connections as the sparse RNN reacts to input patterns multiple times. This plasticity is a form of gated Hebbian synaptic modification; if pattern *μ* is categorized incorrectly, a term 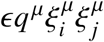 is added to **J**^**s**^, with *ϵ* the learning rate. This makes small corrections to the output of the sparse RNN for incorrectly categorized patterns without significantly interfering with correctly matched patterns. Allowing *ϵ* to decrease with epoch number enhances performance.

Sparse RNNs trained using the OS+ method improve the capacity of OS-initialized networks and maintain the same asymptotic performance [Figure 4OS+]. We find that *α*_OS+ ≈_ 10 *α*_OS_ ≈ 0.01, still 100 times smaller than the performance of BPTT-trained networks. We binned the results and fitted lines as for the OS method and found the ^2^ of the 10 line fits to be 0.993 ± 0.004 and the slope 2.00 ± 0.07. These results were obtained by training sparse RNNs for 150 epochs using the OS+ method. We found that when perfect accuracy was not achieved within 150 epochs, forgetting of previous patterns emerged and accuracy dropped close to chance level within 500 − 1000 epochs.

## Discussion

We presented a biologically plausible neural network architecture that solves the binary classification task with high capacity. This architecture is based on a sparse RNN that solves the task dynamically. Our main purpose was to show that categorization can be achieved in a truly distributed way without the convergence of information onto any single locus. Sparse RNNs can categorize roughly two patterns per plastic synapse, matching classic perceptron performance [3]. These networks are robust to various types of noise and across training methods and activation functions. Our approach supports separate populations of excitatory and inhibitory neurons, and the resulting E/I networks perform well.

The performance of sparse RNNs for categorization, with scaling of the number of patterns proportional to the number of synapses is in keeping with results from other network studies. In sparse Hopfield-type models, the number of stored bits scales with the number of synapses [6]. In Hopfield-style models of recognition memory, the number of patterns than can be identified as familiar or novel scales with the number of synapses [7], a result that also holds for feedforward networks [8] and for various plasticity rules [8, 9].

Categorization by sparse networks has been considered previously by Kushnir and Fusi [2], who used a committee-machine-like readout on a recurrent network with a fixed number of recurrent connections per unit. Fixed here means a number of connections per neuron that was independent of the number of neurons (*N*), as opposed to the case we studied with sparse connections proportional to *N*. In addition, in their study [2], recurrent connections were not learned. Nevertheless, Kushnir and Fusi showed that the recurrent connections play a crucial role in maintaining classification accuracy with sparse readouts. They proved that their model can classify a number of inputs proportional to the number of plastic synapses, as reported by other studies [3, 10, 11] and in ours. Because of the fixed number of connections per neuron, in the regime of large numbers of neurons, their RNN eventually becomes disconnected.

Our results suggest that it is possible to generate decisions in a dynamic and distributed manner in RNNs. This is almost certainly closer than perceptron models to how the bulk of decisions made by biological networks are computed. We suggest the following model: motor or premotor circuits are held in a state of readiness during a go/nogo task, but are not activated until the decision is made. Relevant information is conveyed to the neurons in this circuit, much like the patterns are conveyed to the RNNs we studied. If these inputs are appropriate for a nogo decision, the motor/premotor circuit may be perturbed, but it does not make a transition to a fully activated state. This is analogous to the blue curves in Figure 1C. If, on the other hand, the evidence favors action, the motor/premotor circuit reacts more strongly, analogous to the red curves in Figure 1C, and the motor action is initiated.

## Acknowledgements

Research supported by NSF NeuroNex Award DBI-1707398, the Gatsby Charitable Foundation, and Boehringer Ingelheim Fonds.

## Methods

### BPTT Simulations

We simulated sparse RNNs with discrete time steps according to the dynamics in Equation 1. For all BPTT simulations we used custom RNN architectures and code developed in Python using PyTorch [12]. Code is available at github.com/DenisTurcu/SparseRNN.

### Response function

The shifted, scaled and rectified hyperbolic tangent response function we used for all reported results is max[(tanh(*x*+ *x*_0_) − tanh(*x*_0_))/(1 − tanh(*x*_0_)), 0], where *x*_0_ represents the shift amount. We used *x*_0_ < 0, typically *x*_0_ = −0.5, such that the derivative at small positive values is supralinear.

### Sparsity Mask

We imposed a sparsity mask **M** on the recurrent connections **J** such that the effective recurrent connections **J**^**s**^ ≡ **J** ⊙ **M** are sparse. The sparsity mask **M** is an *N* × *N* matrix with elements equal to either 0 or 1 and Σ_*i,j*,_ **M**_***i j***_ = *f N*^2^. All the 1s are randomly placed in **M**. This ensures that only *f N*^2^ of the *N*^2^ recurrent connections are used by the sparse RNN.

### Decision Threshold

The simulated sparse RNNs are judged based on their output being above or below a certain threshold. For all results reported, except for part of Table 1, we trained the networks using a label-specific dynamic target. The targets for the ± 1 categories [Figure 1C] implicitly define a discrimination threshold 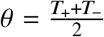. We chose the targets to start at time *t*_ON_ such that the sparse RNNs process their input while it is presented and then make a decision between _ON_ and _F_. The targets are constant at all times *t*_ON_ <*t* < *t*_F_, such that the sparse RNNs are required to maintain their decision for some amount of time and act as evidence accumulators.

### Split E-I Architecture

Excitation-inhibition constraints were imposed at every gradient step during training for all simulations with such restrictions. The constraints ensure that all elements of any given column of **J**^**s**^ have the same pre-assigned sign. We defined a vector **v** of size with elements equal to either 1, for excitatory, or −1, for inhibitory identity. We defined *N*_*E*_ as the number of 1s and *N*_*I*_ as the number of − 1s in **v** such that *N*_*E*_ + *N*_*I*_ = *N*. We updated the effective recurrent connections after each optimization step according to **J**^**s**^_*ij*_ ← **J**^**s**^_*ij*_ sgn (**J**^**s**^_*ij*_)**v**_*j*_, which ensures that all recurrent connections going out from neuron have the same identity as **v**_*j*_.

### Noise Description

No noise was introduced during training the sparse RNNs, yet they are robust to noise at test time. We presented two types of noise after training the sparse RNNs, input noise and dynamic noise. Input noise changes the input pattern presented to sparse RNN. Dynamic noise alters the recurrent state of the sparse RNN at each time step. We incorporated these noise types into Equation 1:

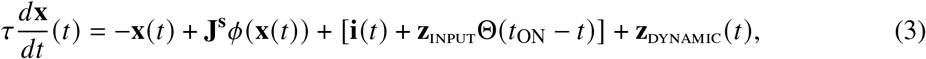

where **z**_INPUT_ is Gaussian, fixed for every trial and turns off with the input at *t*_ON_ as noted by the step-function, and *𝓏*_DYNAMIC_ (*t*) is Gaussian and changes independently at every time step.

### Loss Functions

The primary loss function we used for all reported results, except part of Table 1, was the loss function based on label-specific dynamic targets. This targets-based loss function penalizes the sparse RNNs when their output is below (for + 1 category) or above (for − 1 category) the respective target, in proportion to the absolute value difference between *𝓏 (t)* and *T*_*+(*−)_. To avoid instructing the sparse RNNs how to perform the task via this target-based training, we also trained networks using a threshold-based loss function. For this loss function, we set the discrimination threshold, *θ*, constant in time, similar to before. The threshold-based loss function depends on a nonlinear mapping applied to the quantity *−q*^*μ*^*(𝓏(t) − θ)*^;^and summed from time *t*_F_ −*Δt* to time *t*_F_. This quantity is positive at a time step when there is a categorization miss-match, e.g. *𝓏(t)* < *θ*, but *q*^*μ*^ = +1, and negative otherwise. Therefore, we used a nonlinear mapping which is large for positive values, and small for negative values. Specifically, this mapping was max [*xβ*, 0] −tanh (max [−*xβ*, 0]) for some slope parameter *β*, typically set to 1.

### BPTT Speed Up

BPTT training was sped up with the introduction of “PyTorch’s sparse” module. We started using this module once it reached a stable development version and operations on sparse tensors were properly differentiable. This work replaces our version of using a sparse recurrent connections matrix via a sparsity mask and we used for all results reported, except from [Figure 3D] and Table 1. Code is available at github.com/DenisTurcu/SparseRNN.

### Hebbian Learning

All Hebbian learning results reported were generated using custom Matlab code (Mathworks, Natick, MA). Code is available at github.com/DenisTurcu/SparseRNN

### Directed Graph Percolation

We estimated the probability that a randomly generated directed graph with *N* nodes and connection probability *f* from node *i* to node *j* consists of a strongly connected sub-graph of size *N* − *k* and *k* other nodes empirically. We call this probability 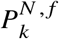. We ran these simulations in Python using Kosaraju’s algorithm and adapted the implementation of the algorithm by Neelam Yadav. Code is available at github.com/DenisTurcu/SparseRNN

